# More than 2,500 coding genes in the human reference gene set still have unsettled status

**DOI:** 10.1101/2024.12.05.626965

**Authors:** Miguel Maquedano, Daniel Cerdán-Vélez, Michael L. Tress

## Abstract

In 2018 we analysed the three main repositories for the human proteome, Ensembl/GENCODE, RefSeq and UniProtKB. They disagreed on the coding status of one of every eight annotated coding genes. The analysis inspired bilateral collaborations between annotation groups.

Here we have repeated our analysis with updated versions of the three reference coding gene sets. Superficially, little appears to have changed. Although there are slightly fewer genes predicted as coding overall, the three groups still disagree on the status of 2,606 annotated genes. However, a comparison without read-through genes and immunoglobulin fragments shows that the three reference sets have merged or reclassified more than 700 genes since the last analysis and that just 0.6% of Ensembl/GENCODE coding genes are not also annotated by the other two reference sets.

We used eight features indicative of non-coding genes to examine the 21,873 coding genes annotated across the three reference sets. We found that more than 2,000 had one or more potential non-coding features. While some of these genes will be protein coding, we believe that most are likely to be non-coding genes or pseudogenes. Our results suggest that annotators still vastly overestimate the number of true coding genes.

## Introduction

Since the initial release of the human genome sequence almost a quarter of a century ago [1,2] annotators have been carrying out a detailed curation of the coding and non-coding genes found in the genome. The analysis of the protein coding gene complement has been implemented principally by three groups of researchers. The Ensembl/GENCODE [3,4] and RefSeq [5] reference sets are based on genomic coordinates, and the UniProtKB [6] proteome is based on the recorded proteins.

New coding genes are added to the reference sets when there is sufficient experimental or conservation evidence to support their annotation, but genes, and their related transcripts and protein isoforms, can also undergo changes in status as existing annotations are revised. Many genes are reclassified as a result of the annotators adjusting to the available evidence, and some of the more difficult to discriminate genes may be reclassified more than once. This has occurred with the pseudogene *WASH6P*, for example [7].

Before the human genome was sequenced, estimates for the number of human coding genes were between 25,000 and 40,000, but since then a number of large-scale analyses have revised the estimates downwards to between 19,000 and 22,000 genes [8–13], with the current number of annotated coding genes in the two coordinate-based reference sets much closer to the lower of the two figures. At present, the number of coding genes in the human genome is increasing somewhat because there is a concerted push to find evidence for small ORFs that might have been missed by the annotators [14].

Annotators often add protein coding genes based on experimental evidence from external research groups. Many research groups have been focused on coding gene discovery, largely because it is easier and more rewarding to find new coding genes than it is to demonstrate that predicted coding genes do not produce proteins. Only a few groups have attempted to predict which coding genes might have been misclassified by annotators [8. 9, 11, 13]. Clamp *et al*. [8] showed that most annotated novel ORFs resembled non-coding RNA rather than coding genes and estimated that there were just 20,500 human coding genes, while Church *et al*. [9] compared the mouse and human gene sets and predicted that the number of human coding genes was below 20,000. Many coding genes were reclassified as not coding after these two analyses.

We have also published two previous analyses in which we attempted to uncover genes that were likely to have been misclassified as coding [11, 13]. We used a series of indicators, called potential non-coding features, to classify genes as potential non-coding [11]. Genes with potential non-coding features had almost no peptide support in large-scale proteomics analyses and many of the potential non-coding genes we identified have been reclassified by manual curators. In 2014, we flagged 2,001 potential non-coding genes in the GENCODE v12 gene set [11]. Almost half of these genes (908) were withdrawn or reclassified from the human reference set by the expert curators in GENCODE. In 2018, using the GENCODE v24 gene set, we labelled another 2,278 genes as potential non-coding [13]. Since then, 379 of these coding genes have been reclassified as not coding.

In the second analysis we also merged the Ensembl/GENCODE, RefSeq and UniProtKB gene sets and found that the union of the three reference sets annotated 22,210 coding genes [13]. We showed that one in eight annotated coding genes were not regarded as coding by all three curation groups [13]. To converge on an agreed set of coding genes, curators of the three reference sets set up joint projects such as MANE [15] after the results of the second analysis.

Here, we have carried out a third analysis of the state of the human coding gene complement. We merged the Ensembl/GENCODE, RefSeq and UniProtKB gene sets once more and found that there has been some convergence between the three databases, but that there are still thousands of potential non-coding genes that are likely to be misclassified as coding by the three curation groups.

## Methods

### The Ensembl/GENCODE reference set

We downloaded the coding genes from the Ensembl 111/GENCODE v45 reference set from BioMart [3], restricting the list to protein coding genes in the 24 distinct human chromosomes and the mitochondrial chromosome. At the same time, we downloaded the equivalent gene/protein identifications that Biomart had stored for UniProtKB and RefSeq. We downloaded 20,444 Ensembl/GENCODE coding genes.

### The UniProtKB reference proteome

We downloaded those reviewed UniProtKB entries that were tagged by UniProtKB as belonging to the UP000005640 (human) proteome in the 27th March 2024 version of the UniProtKB database. Along with the list of proteins we downloaded, we included the HGNC gene names [16] and the Ensembl and RefSeq cross-references that were recorded by UniProtKB. After processing (see below) we ended up with 20,484 entries. UniProtKB have developed an automatic pipeline to annotate Ensembl/GENCODE genes directly into UniProtKB Trembl. Many of these are included in the UniProtKB human proteome, but we did not include these unreviewed entries in our analysis. More than two thirds of the UniProtKB Trembl sequences we left out were Immunoglobulin/T-cell receptor fragments or from read-through genes.

### The RefSeq reference gene set

We downloaded the human protein coding genes from the NCBI gene database. This produced 20,441 coding genes. We selected only those in the GRCh38 assembly because RefSeq has already includes predictions for the CHM13 assembly. Of these, seven were coding loci that covered the immunoglobulin and T-cell receptor fragment clusters. Individual immunoglobulin and T-cell receptor fragments are defined as “OTHER” in the RefSeq genome. The final list of coding genes was 19,950.

### Merging the three sets

The sets were merged initially via their HGNC symbols and discrepancies were curated manually. Since the UniProtKB database is based on proteins rather than genes, UniProtKB had the most obvious discrepancies. Some genes had multiple UniProtKB entries (e.g. *TMPO* which has entries for alternatively spliced proteins LAP2 alpha and LAP2 beta/gamma), while some UniProtKB protein-sequence identical entries pointed to multiple genes (e.g. entry Q9ULZ0 which maps to six “coding” genes, *TP53TG3*, *TP53TG3B*, *TP53TG3C*, *TP53TG3D*, *TP53TG3E* and *TP53TG3F*). Both sets of discrepancies had to be de-duplicated so that each coding gene corresponded with a single UniProtKB entry. In addition, UniProtKB includes proteins that came from genes only present in scaffolds. These were removed.

Other corrections were made based on BioMart cross-references and UniProtKB ID mapping cross-references where necessary. In more complicated cases, we checked the coordinates of the Ensembl/GENCODE and RefSeq genes with the UCSC genome browser [17]. RefSeq and Ensembl did have different numbers of coding genes at the same locus in a small number of cases. For example, the gene *ERCC6*, where Ensembl/GENCODE has a single gene and RefSeq two. In these cases, we merged one of the multiple genes and left the second one without a match in the other reference set.

### Determining coding status of non-intersection genes

For the 2,606 coding genes outside of the intersection between the three sets (genes annotated as coding in just one or two of the three reference sets, but not in all three), we determined the status of the gene in the reference set (or sets) in which it was not annotated as coding. The most common gene status were read-through gene (669), pseudogene (484), IG/TR fragment (429), antisense (239), lncRNA (217), intergenic (114), UTR ORF (92), intronic (84), and retroviral (72).

Read-through genes are tagged by GENCODE [4], and the status of pseudogene was applied if the coding genes overlapped a pseudogene in the other reference set, or if the gene had an HGNC name that indicated that the entry was a pseudogene. The IG/TR fragment gene status was applied to those genes that were tagged as immunoglobulin or T-cell receptor fragments in Ensembl/GENCODE, and to those that were clearly immunoglobulin or T-cell receptor fragments from the associated HGNC gene name.

Antisense status was applied when transcripts were on the opposite strand of a coding gene or a pseudogene and close to or overlapping the UTR of those genes, or when the HGNC name indicated that it was antisense to another gene. Genes were labelled as lncRNA when they overlapped exons from lncRNA in the other reference set or if the gene had an HGNC name that indicated that the entry was originally from a lncRNA gene.

The status “intergenic” was reserved for those genes that did not overlap any other type of gene in the other reference set, while those tagged as “no coordinates” were all from UniProtKB only did not map to the GRCh38 genome. UTR ORF covered those coding genes that overlapped with UTR in the other reference set, intronic covered those coding genes were found wholly within introns of genes in the other reference set, and artefact and TEC covered those coding genes that were annotated as such in the equivalent in the Ensembl/GENCODE reference set. Genes were labelled as retroviral when they overlapped exons from retroviral genes in the other reference set or if the gene had an HGNC name that indicated that the entry was originally from a retrovirus.

### Potential non-coding features

Potential non-coding features were taken directly from the UniProtKB and/or the Ensembl/GENCODE annotations. The three exceptions were PhyloCSFMax, which is the standard PhyloCSF [18] score of the highest scoring exon that has a minimum 10 codons; gene family age, which is the normalised age of the most distant BLAST [19] hit from an APPRIS database [20] search against a limited list of RefSeq reference proteomes [5]; and “No protein features” which applies to those genes that do not have functional residues, homology to protein structure or domains, cross-species conservation or trans-membrane helices (all data from APPRIS modules), and in addition have a negative PhyloCSFMax score. Cut-offs used for potential non-coding features were −14 or lower using PhyloCSFMax were and no homologues beyond catarrhini for APPRIS gene age.

### Peptide data

Peptide data is available from the APPRIS database for all Ensembl/GENCODE genes. We also downloaded peptides from PeptideAtlas [21]. In the case of PeptideAtlas, all peptides had to be tryptic peptides and have at least two observations [21]. Peptides that mapped to more than one gene were left out of the analysis. Peptides were not filtered further and we did not check for false positives. In this analysis we did not use the peptides as validation for coding genes or as a potential non-coding feature, merely as an indicator.

### Non-synonymous to synonymous ratios

To evaluate the ratio of non-synonymous to synonymous variants (NS/Syn ratio) as an indicator of potentially non-coding genes, we first processed variant data from GNOMAD genomes analysis [22] using the Variant Effect Predictor (VEP) tool [23]. We utilised the principal transcripts derived from the APPRIS database as the representative for each gene to maximise the coding potential [24].

Following this, the VEP output was parsed into a format suitable for downstream analyses. Variants were filtered to include only those containing one of the four valid alleles (A, C, T, G). The dataset was then further refined by retaining only the following types of variants: ‘start_lost,’ ‘synonymous_variant,’ ‘missense_variant,’ ‘stop_gained,’ ‘stop_lost,’ ‘splice_donor_variant,’ and ‘splice_acceptor_variant.’ Additionally, ‘splice_region_variant’ was included if it co-occurred with other valid variant types, as these are counted according to their accompanying variant classification.

Frameshift variants were excluded from the analysis. We also removed variants lacking a Frequency_maxAF annotation (i.e., not included in the gnomAD database), and those with a maxAF of 0. Variants with a maxAF > 0.005 were considered rare for the posterior analyses. To assess the NS/Syn ratios across rare and common alleles in different subsets of potentially non-coding genes we used just the missense and synonymous variants.

Potential non-coding features were defined from these NS/Syn ratios. If the NS/Syn ratio for the subset of genes was higher than 2.0 in both common and rare alleles and the difference between the two sets was less than 0.3, the feature was determined to be potential non-coding. We defined 7 potential non-coding features this way and added all read-through genes as an eighth potential non-coding feature. These genes appear to be partly under purifying selection because most of their exons overlap known coding exons.

## Results

The merge of the three reference sets found 21,873 coding genes, slightly fewer than the 22,210 we counted in the previous merge [13]. The distribution of coding genes between the three reference sets in the two merges can be seen in figures 1A and 1B. Of the 21,873 coding genes, 19,267 genes are predicted to be coding by all three sets of annotators, which means there were 179 fewer genes agreed upon by all three databases with respect to our previous analysis. The three reference annotations fail to agree on the coding status of 2,606 genes, so close to one in eight annotated coding genes still have different status in at least one of the three reference sets.

**Figure 1.**
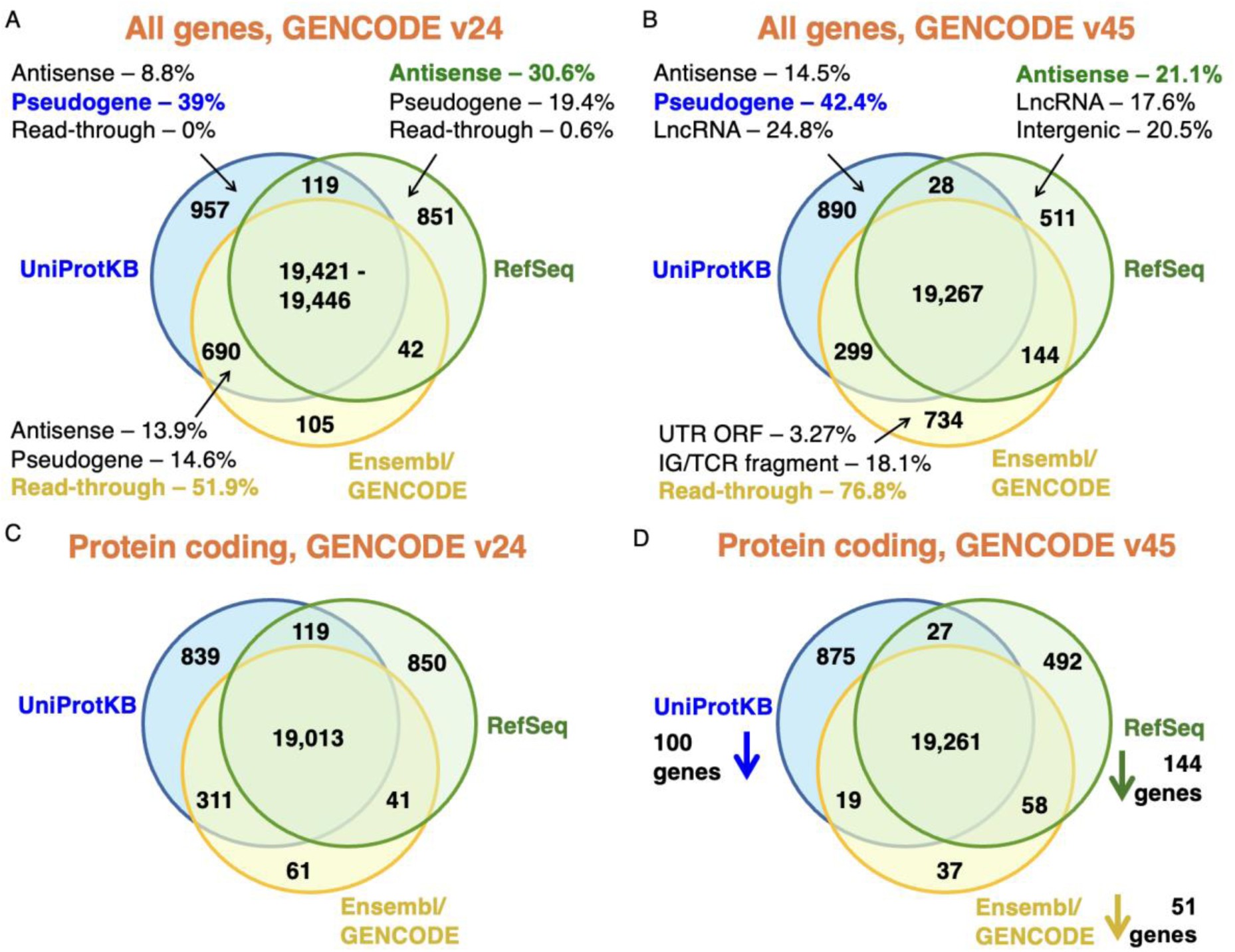
Reference coding genes contemporary to the GENCODE v24 and v45 sets. Coding genes in the merged sets of UniProtKB, RefSeq and Ensembl/GENCODE (see methods). In A, the count of all genes classified as coding in each of the three reference databases and the intersection between them at the time of release of the GENCODE v24 reference set [13]. In B, the count of all genes classified as coding in each of the three reference databases coinciding with the release of the GENCODE v45 reference set. Panel C shows the counts for the merge of the three annotations contemporary to the GENCODE v24 reference set excluding all read-through genes and immunoglobulin and T-cell receptor fragments. In panel D, the same count, but at the time of the GENCODE v45 reference set.

Read-through genes, a class of genes in which all transcripts are read-through transcripts, make up a quarter (25.7%) of the genes that have different status in at least one of the three reference sets. Almost one in five genes with disputed coding status (18.6%) are annotated as pseudogenes by at least one of the other databases. For coding genes annotated by just one of the three databases we have detailed the most common reasons for the discrepancies with the other two reference sets (figures 1A and 1B). More than 40% of UniProtKB unique coding genes are regarded as pseudogenes by the other two annotation groups, three quarters of Ensembl/GENCODE unique genes are read-through genes and many RefSeq unique genes are likely non-coding with antisense and LncRNA labels making up more than a third of differences.

### Merging the three reference sets without read-throughs or immunoglobulin fragments

A sixth of the coding genes not agreed on by all gene annotators were immunoglobulin or T-cell receptor fragments. Even though T-cell receptor and immunoglobulin fragments produce protein regions, they are not always considered to be true protein coding genes, and in fact RefSeq annotates them as “other” rather than coding and annotates seven protein coding loci in their place. Much of the decrease in RefSeq gene numbers since 2018 is down to this technicality. Many genes moved from the intersection in 2018 to the Ensembl/GENCODE and UniProtKB set in this analysis.

In the previous analysis we include all the UniProtKB human proteome entries, including the non-reviewed entries. This time we left out the unreviewed sequences out of the merge. The unreviewed entries were almost entirely proteins from read-through genes and T-cell receptor and immunoglobulin fragments, which pushed these genes out of the agreement between UniProtKB and GENCODE in the 2018 analysis and into the Ensembl/GENCODE only category in this analysis (figures 1A and 1B).

Given that immunoglobulin and T-cell receptor fragments and many read-through genes have moved from one set to another in this analysis, we carried out a comparison between the two merges leaving out the immunoglobulin and T-cell receptor fragments and read-through genes. Removing these genes from the comparison ought to provide a much better perspective of the effects of the Ensembl/GENCODE bilateral collaborations on the three reference sets.

When T-cell receptor and immunoglobulin fragments and read-through genes are set aside from the comparison with the 2018 analysis [13], it becomes clear that all three reference sets now annotate substantially fewer coding genes (figures 1C and 1D). At the same time they agree on 248 more genes than they did in 2018. In fact, 713 of the coding genes that were outside of the intersection have been merged or reclassified since the 2018 analysis, which means 32.1% fewer disagreements between the three sets.

When we leave out T-cell receptor and immunoglobulin fragments and read-through genes from the merge, almost all of the coding genes annotated by Ensembl/GENCODE (99.4%) are in agreement with the other two databases. If read-through genes had been reclassified as not coding, Ensembl/GENCODE would have had just 114 coding genes that were not part of the other two reference sets (figures 1C and 1D). The merge without read-through genes is not just a theoretical exercise, the Ensembl/GENCODE curators plan to reclassify read-through genes as not coding. When they do this, many of the disagreements with the other two databases will disappear.

### Potential non-coding features for GENCODE45 genes

In our previous analysis of the state of the human proteome [13], we defined 16 potential non-coding features. These were drawn from the UniProtKB and Ensembl annotations and from GENCODE consortium tools [18, 20]. We used these potential non-coding features to tag 2,278 Ensembl/GENCODE genes as potential non-coding. Coding genes with one or more of these 16 features had very few peptides in large-scale tissue-based proteomics experiments [13]. They also had significantly lower transcript expression, many more copy number variants, twice as many potential high impact variants and much higher synonymous to non-synonymous rates than other coding genes [13]. We theorised that many of these potential non-coding genes may not be *bona fide* coding genes.

For this paper, we used non-synonymous to synonymous (NS/Syn) ratios to select potential non-coding features. We classed a feature as potential non-coding if the difference between the common and rare NS/Syn ratios for genes with the feature was smaller than 0.3, and if the common and rare NS/Syn ratios were both above 2.0. As a comparison, coding genes with peptide support have a rare NS/Syn ratio of 1.96 and a common NS/Syn ratio of 0.9, so are clearly under selection pressure.

Just seven features that met these criteria for protein coding genes (figure 2), UniProtKB cautions (“dubious CDS” and “possible pseudogene”), UniProtKB “uncertain” evidence code, the word “pseudogene” in the UniProtKB protein description, recent gene family age (calculated through the APPRIS database [20]), poor PhyloCSF conservation [18], lack of protein features (calculated from APPRIS and PhyloCSF scores) and the HGNC gene name [16]. Four of these features were maintained from the previous analysis, two are modified and one is new. The PNC features from the previous analysis that were excluded either tag minimal numbers of genes or have NS/Syn ratios that did not meet the criteria for this paper. Two of the main potential non-coding features from the previous analysis, UniProtKB “homology” and “predicted” evidence codes had high NS/Syn ratios over common alleles, but these ratios were more than 0.3 points lower than the equivalent NS/Syn ratios for rare alleles (supplementary figure 1). Not including these two features as potential non-coding is the main reason why there are many fewer potential non-coding genes in this analysis.

**Figure 2.**
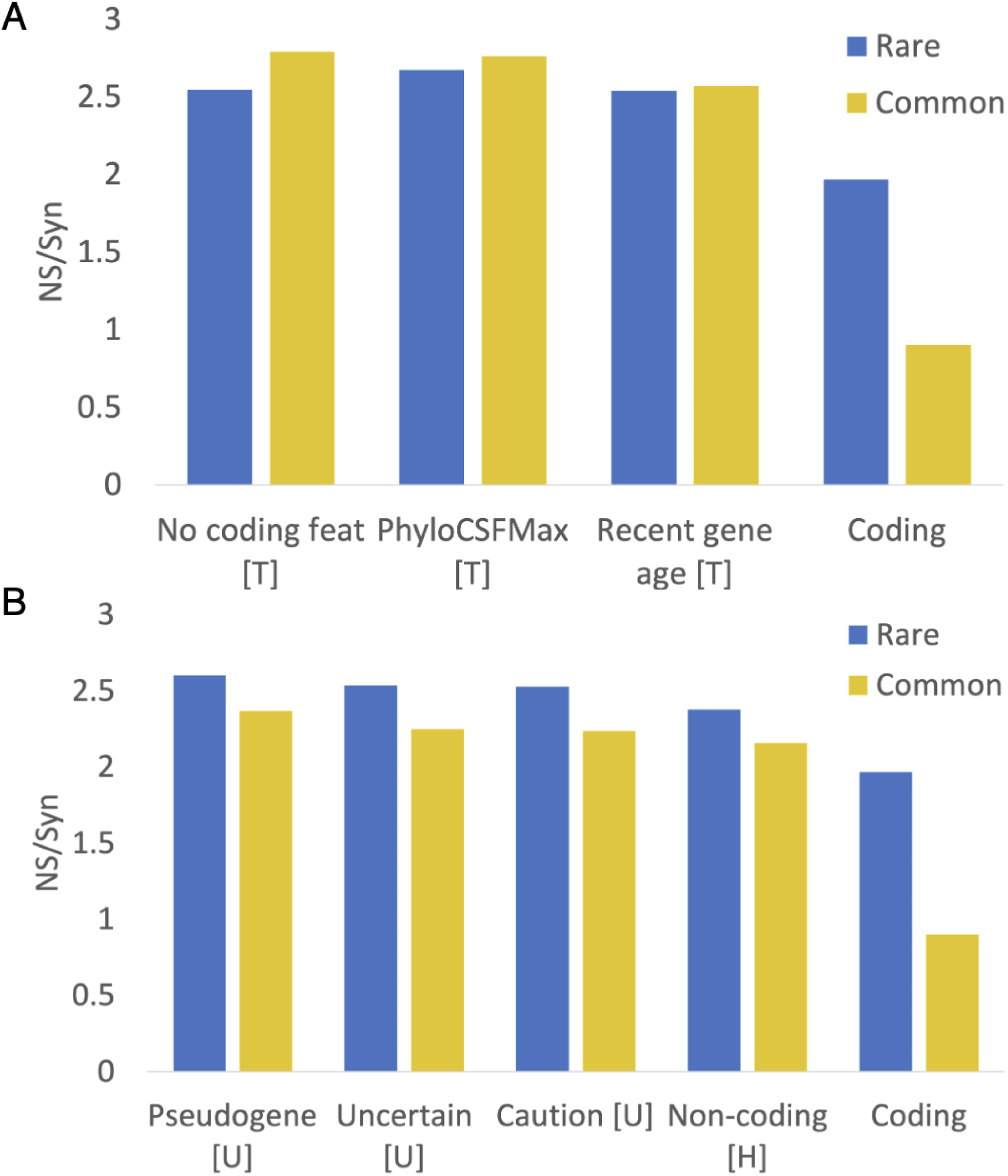
NS/Syn ratios for potential non-coding features in Ensembl/GENCODE. Panel A shows the calculated NS/Syn ratios for the three potential non-coding features that are based on a lack of protein-like conservation and are calculated with GENCODE tools PhyloCSF and APPRIS. These features, marked with a “T”, were calculated in house from GENCODE data and are detailed in the methods section. Panel B shows NS/Syn ratios for features taken directly from annotation by manual curators. Features with a “U” came directly from the UniProtKB database. Features marked with an “H” came directly from the HGNC name. In both cases the NS/Syn ratios for coding genes supported by PeptideAtlas peptides are included as a control.

We added read-through genes to the final list of potential non-coding features even though the common variant NS/Syn ratio of this feature was below the threshold. Read-through genes will inevitably have lower common variant NS/Syn ratios because a large majority of their exons overlap coding exons in the same frame. The final list of potential non-coding features is shown in table 1. With these eight potential non-coding features we tagged 1,118 genes in GENCODE v45 (5.5%) as potential non-coding.

**Table 1.** The eight potential non-coding (PNC) features, the number of genes tagged by each feature and the number of those genes that had at least two supporting peptides in the PeptideAtlas database. The letter in brackets refers to the source of the feature, GENCODE tools APPRIS and PhyloCSF (T), the GENCODE annotation (G), UniProtKB (U), and HGNC (H). Read-through genes were left out of the proteomics analysis to avoid moonlighting peptides that mapped to more than one gene.

| **PNC feature** | **Genes** | **Peptide support** |
| --- | --- | --- |
| *Recent gene age [T]* | 241 | 19 |
| *No protein features [T]* | 118 | 20 |
| *Read-through gene [G]* | 655 | 0 |
| *PhyloCSF maximum [T]* | 98 | 10 |
| *Caution note [U]* | 91 | 24 |
| *Uncertain evidence [U]* | 83 | 22 |
| *Pseudogene [U]* | 31 | 16 |
| *Non-coding name [H]* | 38 | 5 |

### Protein evidence for non-coding GENCODE v45 potential non-coding genes

Peptide support was calculated for each gene from the peptides in the PeptideAtlas database [21]. Table 1 shows the number of genes with each potential non-coding feature that are supported by at least two tryptic peptides in PeptideAtlas. PeptideAtlas has peptide data from 3,160 mass spectrometry experiments, so it has a much higher coverage than the experiments we used in our previous analysis [13]. PeptideAtlas provides peptide support for 89.2% of the GENCODE v45 genes not tagged as potential non-coding. The disadvantage of using such a large source of peptide support is that we cannot validate the spectra, so a proportion of the identifications are almost certainly false positives. Given the similarity to known coding genes, the features that are most likely to have false positive matches are those that involve pseudogenes. It is not surprising that one of these has the highest proportion of peptide support (UniProtKB pseudogene, 51.6%). Genes with the least evidence of cross-species conservation (recent gene age, 7.9%, and poor PhyloCSF, 10.2%) have the least peptide support.

### Potential non-coding genes outside of GENCODE

Over all three curated sets, 2,056 of the 21,873 coding genes were tagged as potential non-coding (9.4%). Compared to the genes in the intersection, a much higher proportion of genes that had conflicting status were tagged as potential non-coding (figure 3); 1,781 of genes that were not annotated as coding in all three reference sets (68.3%) had PNC features, against just 275 with potential non-coding features among genes annotated in all three databases (1.4%).

**Figure 3.**
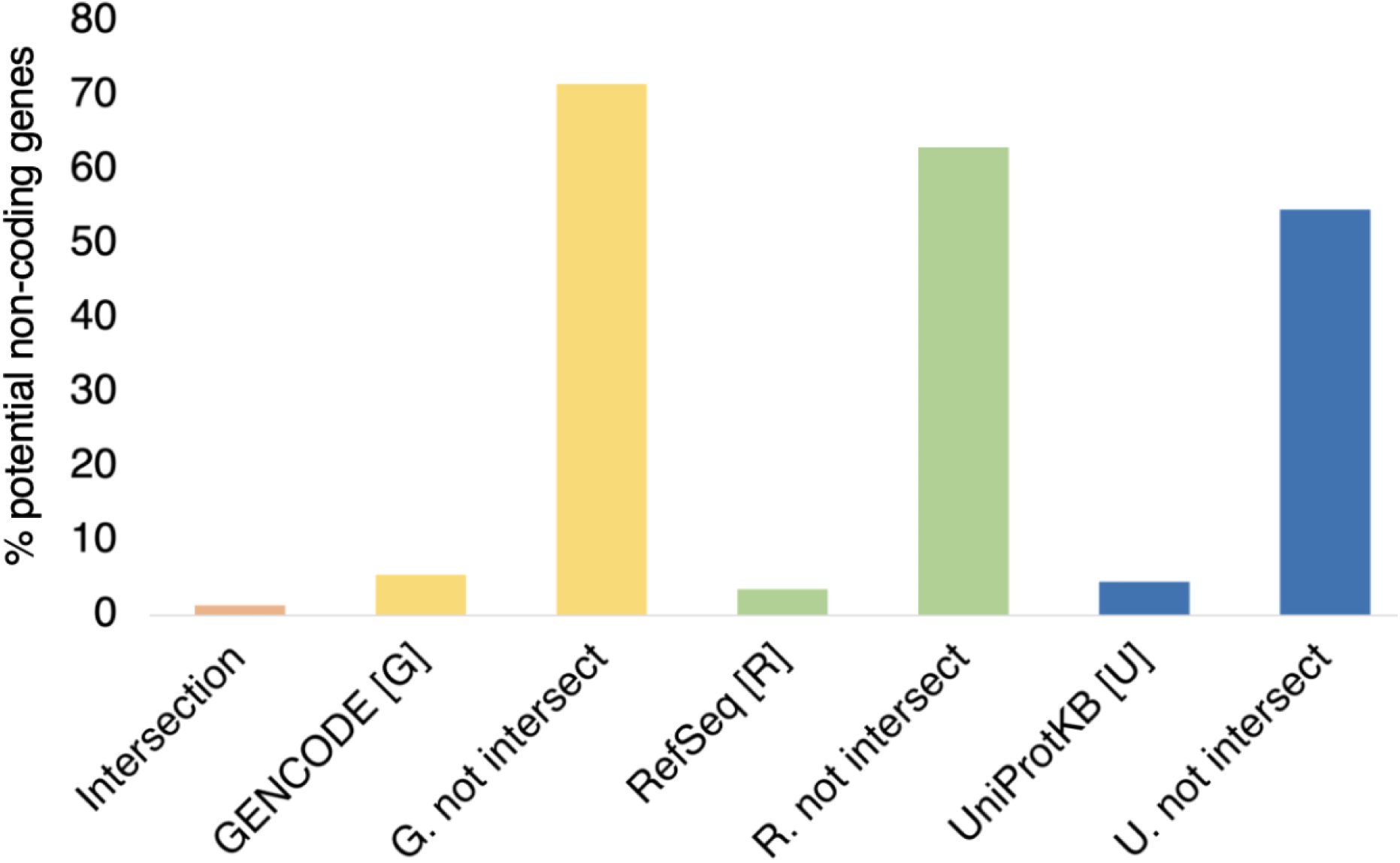
Potential non-coding genes inside and outside of the intersection. The percentage of the genes in different gene sets that are tagged as potential non-coding genes. The genes in each set are those shown in figure 1B, “GENCODE [G]” are all Ensembl/GENCODE genes, “RefSeq [R]” are all RefSeq genes, and “UniProtKB [U]” are all UniProtKB genes. “G. not intersect” are all Ensembl/GENCODE genes that are not in the intersection between the three sets. “R. not intersect” are all RefSeq genes that are not in the intersection between the three sets. “U. not intersect” are all UniProtKB genes that are not in the intersection between the three sets.

Even though the potential non-coding features were designed for Ensembl/GENCODE genes, the proportion of UniProtKB proteins outside of the intersection that were predicted to be translated from potential non-coding genes was also high, 666 of 1218 (54.7%). Finally, even though only three of the potential non-coding features could be used for the RefSeq genes, almost two thirds of the RefSeq genes not in the intersection (63.1%) were tagged with potential non-coding features. In fact, the 683 RefSeq genes outside of the intersection were clearly not under purifying selection; over these genes the NS/Syn ratio for rare alleles was 2.32, while the NS/Syn ratio for common alleles was 2.28 (supplementary figure 2).

### Recently evolved potential non-coding genes

Many of these potential non-coding genes in the intersection are clearly coding despite the potential non-coding features. For example, *MGAM2* and *SRGAP2C* have more than 50 peptides each in PeptideAtlas. They are only tagged as potential non-coding because UniProtKB still includes the word “pseudogene” in the description. When UniProtKB updates its annotations, these genes will no longer be potential non-coding. However, there are numerous potential non-coding genes in the intersection that are just as clearly not coding. For instance, there are 6 read-through genes that are annotated as coding in all three reference sets, and none of these are *bona fide* protein coding genes.

There are other clear examples of coding genes with little or no supporting information besides the read-through genes. *HEPN1* is one, a predicted coding gene that is antisense to *HEPACAM* on chromosome 11. *HEPACAM* was first found in liver [25] but is predominantly expressed in the brain. Curiously, *HEPN1* was identified by the same group before they described *HEPACAM* [26], but at no point was it established that *HEPN1* had a protein product. No peptides are detected for *HEPN1* in PeptideAtlas [21] and the Human Protein Atlas [27] antibody identification is tagged as uncertain.

The predicted coding frame of *HEPN1* is not conserved beyond great apes because of a frameshift mutation. Meanwhile, the eight most common GNOMAD germline variants in *HEPN1* are non-synonymous or high impact [22]. Most interestingly, *HEPN1* has an almost identical expression pattern to *HEPACAM* (figure 4A). The evidence suggests that *HEPN1* should be classed as a natural antisense transcript of *HEPACAM* [28]. It may be involved in regulation but is almost certainly not a coding gene.

**Figure 4.**
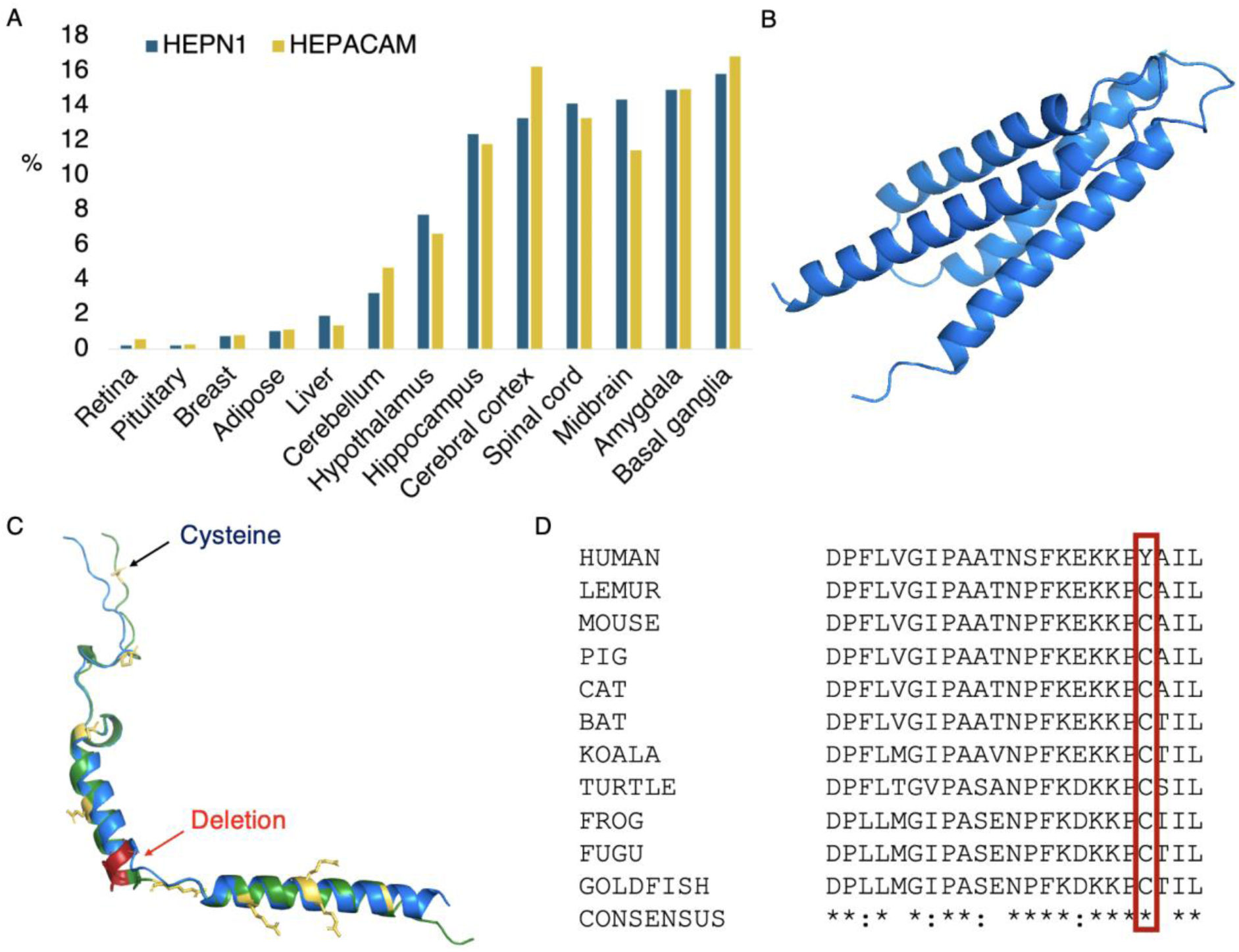
Potential non-coding genes *HEPN1*, *PBOV1* and *GNG14*. Panel A shows the Human Protein Atlas transcript expression per tissue as a percentage of all the detected expression. *HEPN1* (blue) and *HEPACAM* (yellow) have almost identical expression profiles. Panel B shows the predicted AlphaFold [29] structure for *PBOV1*. Panel C superimposes predicted AlphaFold structures of human (blue) and cat (green) *GNG14*. Residues in conserved positions that have mutated in the human sequence are shown as yellow sticks; the deleted human region is mapped onto the cat *GNG14* structure in red. In panel D, an alignment of the C-terminal of vertebrate *GNG14* sequences. The human-specific tyrosine for cysteine swap is marked with a red box.

*BLID* is another single exon gene discovered on the same chromosome and at more or less the same time as *HEPN1* [30]. Its identification as a protein coding gene was partly based on an imaginary BH3-like motif. *BLID* would be a human-specific protein coding gene because only the human ORF has an ATG. Outside of great apes the ORF is broken by frame shifts. Although there are papers that ascribe functions including binding to *BCL2L1* [31] and regulation of the Akt pathway [32], it seems unlikely that the Akt pathway - conserved across eukaryotes - would be controlled by a human-specific gene. As would be expected if this gene were not a coding gene, there is no evidence of purifying selection from germline variants [22], no peptides are detected for *BLID* in PeptideAtlas, and none of the Human Protein Atlas [27], FANTOM [33] or GTex [34] RNA datasets have transcription evidence for *BLID*, not even in breast tissue where it is supposed to be highly expressed [35]. Human Protein Atlas does record minimal transcript evidence for a small number of cancer cell lines. This lack of experimental evidence does suggest that *BLID* is most likely to be a non-coding cancer antigen. As a curiosity, one further feature that *BLID* has in common with *HEPN1* is that both have retracted papers that support functional roles [36, 37].

*PBOV1* is also a single exon gene. There are papers with western blots that support the *PBOV1* protein in prostate and ovarian cancers [38, 39] and RNA support in lymphoma and leukemia in the Human Protein Atlas. AlphaFold [29] predicts that any *PBOV1* protein would fold into a 4-helix bundle (figure 4B), though the prediction is low quality. However, *PBOV1* would be another human-specific coding gene because all primate species have frame shifts relative to human. The eight most common GNOMAD germline variants are non-synonymous or high impact, so there is no hint of purifying selection to support any *de novo* function. As with *BLID*, there are no peptides detected for *PBOV1* in PeptideAtlas and there is no protein or transcript support in normal tissues in Human Protein Atlas. Again, like *BLID*, *PBOV1* appears to be a cancer antigen rather than a standard coding gene. There are plenty of other potential non-coding genes that are likely cancer antigens, including *HMHB1*, *DCANP1* and *PRAC2*. There are also a number of known cancer antigen families. Although cancer antigens were once thought to be coding, most are now thought to be aberrant [40].

### Potential non-coding genes and pseudogenization

While all these genes have appeared recently, there are also genes with evolutionary history but that do not appear to be coding. One such case is *GNG14*, a two-exon gene that is clearly coding in most mammalian species. *GNG14* would produce a heterotrimeric G protein gamma subunit, one of three proteins in a GTP binding complex that mediates signal transduction across the plasma membrane [41]. There 13 other G protein gamma subunit isoforms annotated in the human genome.

The coding exons of the other G protein gamma subunit isoforms are highly conserved across mammals, but *GNG14* seems to have lost importance among monkeys because about 20% of species have ORF-disabling mutations. Great apes in particular have three radical mutations in positions that are highly conserved for all GNG proteins. Human *GNG14* has a frameshift mutation close to the 5’ splice site of the second exon that can be skipped, but only at the cost of eliminating three of the most conserved amino acids in *GNG14* (Figure 4C).

It is known that G protein gamma isoforms are post-translationally modified at the C-terminus. The cysteine residue four amino acids from the C-terminus is vital to the function of the G protein gamma isoforms because it is prenylated, the last three amino acids are removed, and finally the prenylated cysteine is carboxymethylated [42]. The carboxymethylation is required for the association to the plasma membrane, and without it the complex cannot bind receptors or interact with downstream effector proteins [43, 44]. In great apes, this cysteine would be mutated to tyrosine, so the GNG14 isoform function is abolished (figure 4D).

If there were any human *GNG14* protein, it would clearly not function as a G protein gamma isoform. There is also no evidence that the mutated isoform might be functionally important in any other role either - two of the 3 most common germline variants in GNOMAD are a stop gain and a start lost variant. There is also no protein or transcript evidence for *GNG14*, and it has no supporting publications. *GNG14* is quite clearly a misannotated pseudogene in great apes, and possibly also in monkeys.

Connexins are four trans-membrane helix bundles that associate to form gap junction channels that allow electrical signal transfer and metabolite diffusion between cells [45]. There are more than 20 gap junction isoforms annotated in the human genome. One of these, *GJE1*, has just four extra-cellular cysteines instead of the usual six, and mouse experiments have shown that it does not form gap junctions [46]. *GJE1* is clearly conserved across vertebrates, but conservation evidence shows just as clearly that *GJE1* is a pseudogene across primates. Human, chimp and gorilla *GJE1* transcripts do still have intact ORFs, though gorilla has lost one of the four remaining conserved extracellular cysteines. There is no supporting experimental evidence for *GJE1* in PeptideAtlas or the Human Protein Atlas.

Given that *GJE1* probably pseudogenized at least 80 million years ago, it raises the question, which is more unlikely, that the human and chimp ORFs maintained their frames over 80 million years despite being pseudogenes, or that the chimpanzee and human ancestor were the only two species that escaped pseudogenization, yet there is no physical evidence for their expression? Fortunately, there are clues that can be used to come to a decision. Firstly, the human, chimpanzee and gorilla have radical amino acid changes [47] in conserved positions that will almost adversely affect the packing of the trans-membrane helices (figure 5A). Secondly, the 5’ end of the third coding exon of *GJE1* in human, chimpanzee and gorilla has had to be shortened by four codons by the annotators in order to escape a stop codon that is unique to these three species.

**Figure 5.**
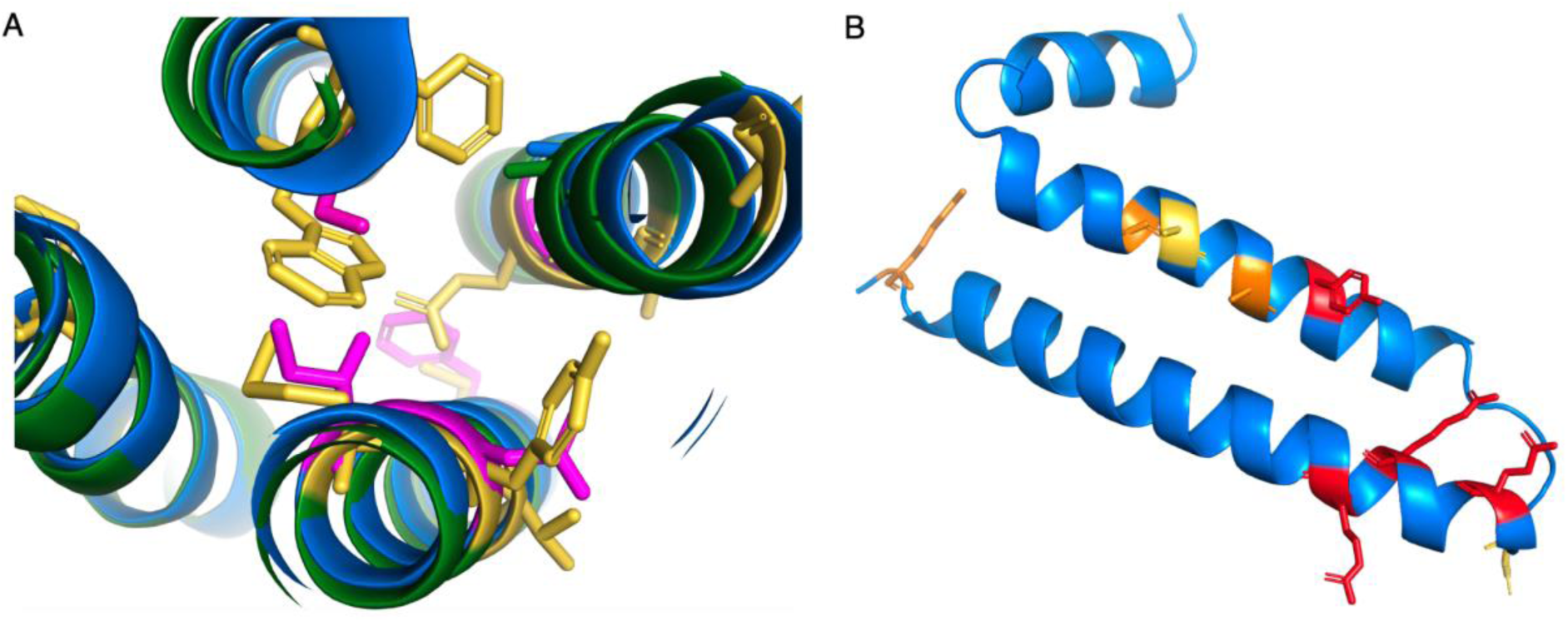
Potential non-coding genes *GJE1* and *HIGD2B*. Panel A shows a cross-section of the predicted AlphaFold structures of human (blue) and cat (green) *GJE1*. Cat *GJE1* conserved residues that have mutated in the human sequence are shown as yellow sticks; some of the mutated human residues are shown magenta. Cat *GJE1* forms a four-helix trans-membrane bundle, but many of the human mutations would adversely affect the packing of the transmembrane helices. From top, clockwise, tryptophan (cat) to serine (human), glutamate to alanine, tyrosine to asparagine, methionine to isoleucine, and in the background, cysteine to phenylalanine. The tryptophan to serine mutation would require the human sequence to use the fictitious 5’ splice site. In panel B, the predicted AlphaFold structure of human *HIGD2A*. Residues in conserved positions that have mutated in the human *HIGD2B* sequence are shown as sticks, sticks are coloured red when the mutation is in a wholly conserved position and orange when the mutation is in an almost completely conserved position, and yellow when it is another radically different amino acid in a less conserved position.

To make up for the loss of four amino acids in the protein from the shortened 5’ end of the third coding exon, the 3’ end of the second coding exon has been extended by three codons downstream. This mutation is possible, and might produce a functional protein without too much loss of function since the swap occurs at the end of an exon, but there is no experimental evidence for the novel splice site (the cDNA from RefSeq is an inferred model), and the novel 5’ intronic splice acceptor site in exon 2 required to explain the gene models of human, chimpanzee and gorilla *GJE1* would be GTGTGC, which is not an accepted splice acceptor pattern [48]. The predicted structural model suggests that the swap would adversely affect the packing between two transmembrane helices (figure 5A).

The stop codon in the final exon, the radical changing that would affect the packing of the trans-membrane helices (figure 5A) and the lack of experimental support for the inferred novel splice site is compelling evidence that *GJE1* is just as much of a pseudogene in human, chimpanzee and gorilla as it is for all other primate species.

One final example of potential non-coding genes annotated by all three reference databases is the gene *HIGD2B*. *HIGD2B* was identified by both Clamp *et al* [8] and Church *et al* [9] as a pseudogene and has been tagged as a potential non-coding gene in all three of our analyses [11, 13]. It duplicated from *HIGD2A*, an ancient, conserved gene involved in the stabilization of the cytochrome C oxidase (complex IV) in the mitochondrial membrane [49], via retrotransposition at the base of catarrhini. The loss of *HIGD2A* has been shown to result in a damaged complex IV that lacks *COX3* [50]. *HIGD2A* has widespread expression, *HIGD2B* is expressed only in testis.

*HIGD2B* is annotated as coding because the ORF is intact and because there is plenty of transcript evidence in testis. Despite that, and despite the large number of testis tissue experiments in PeptideAtlas, there is no supporting protein evidence at all. Although *HIGD2B* has undergone substantial changes in sequence since retrotransposition and could conceivably have gained a new function, this novel function was obviously not important enough to stop it becoming a pseudogene across all non-ape old world monkeys and in orangutans.

Many of the amino acid changes in human *HIGD2B* are radical [47]. We generated an alignment of 35 vertebrate *HIGD2A* proteins that has 29 completely conserved amino acid positions (supplementary figure 3) and four of these positions, along with another 3 positions that are almost completely conserved, have different amino acids in *HIGD2B* proteins (figure 4B). Three of these changes would be radical. In addition, gorilla *HIGD2B* has two more radical amino acid changes and gibbon *HIGD2B* has another three. The changes are such in gorilla that AlphaFold [29] predicts that the second membrane spanning helix will be four residues shorter.

These radical changes, the widespread pseudogenization of *HIGD2B* across monkeys, and a complete lack of protein evidence in all tissues (including testis) in PeptideAtlas and Human Protein Atlas, strongly suggest that *HIGD2B* is a pseudogene that has not yet been affected by an ORF-breaking mutation.

### Some non-flagged genes are also likely to be non-coding

There are also genes in the intersection of the three reference sets that are not tagged as potential non-coding, but that are highly likely to be misclassified as coding. We have already detailed two genes in previous papers, neither *WASHC1* [7] nor *PLK5* [13] are tagged as potential non-coding in this analysis. Another obvious case and not tagged as potential non-coding is *FTCDNL1*, an uncharacterised five exon gene related to osteoporosis [51]. It is an ancient gene with clear coding conservation across mammals, but conservation across primates is patchy (figure 6A). It seems to be conserved in lemurs but is clearly no longer coding in new world monkeys; in marmosets, for example, all five coding exons have high impact mutations. Among old world monkeys, it is less obvious whether *FTCDNL1* is still a coding gene, but in humans, chimpanzees and gorillas, it has almost certainly become a pseudogene because all four species have lost the final coding exon. In addition to the lost exon and stop codon, gorilla *FTCDNL1* has a unique frameshift mutation and human *FTCDNL1* has a unique stop codon in the third exon.

**Figure 6.**
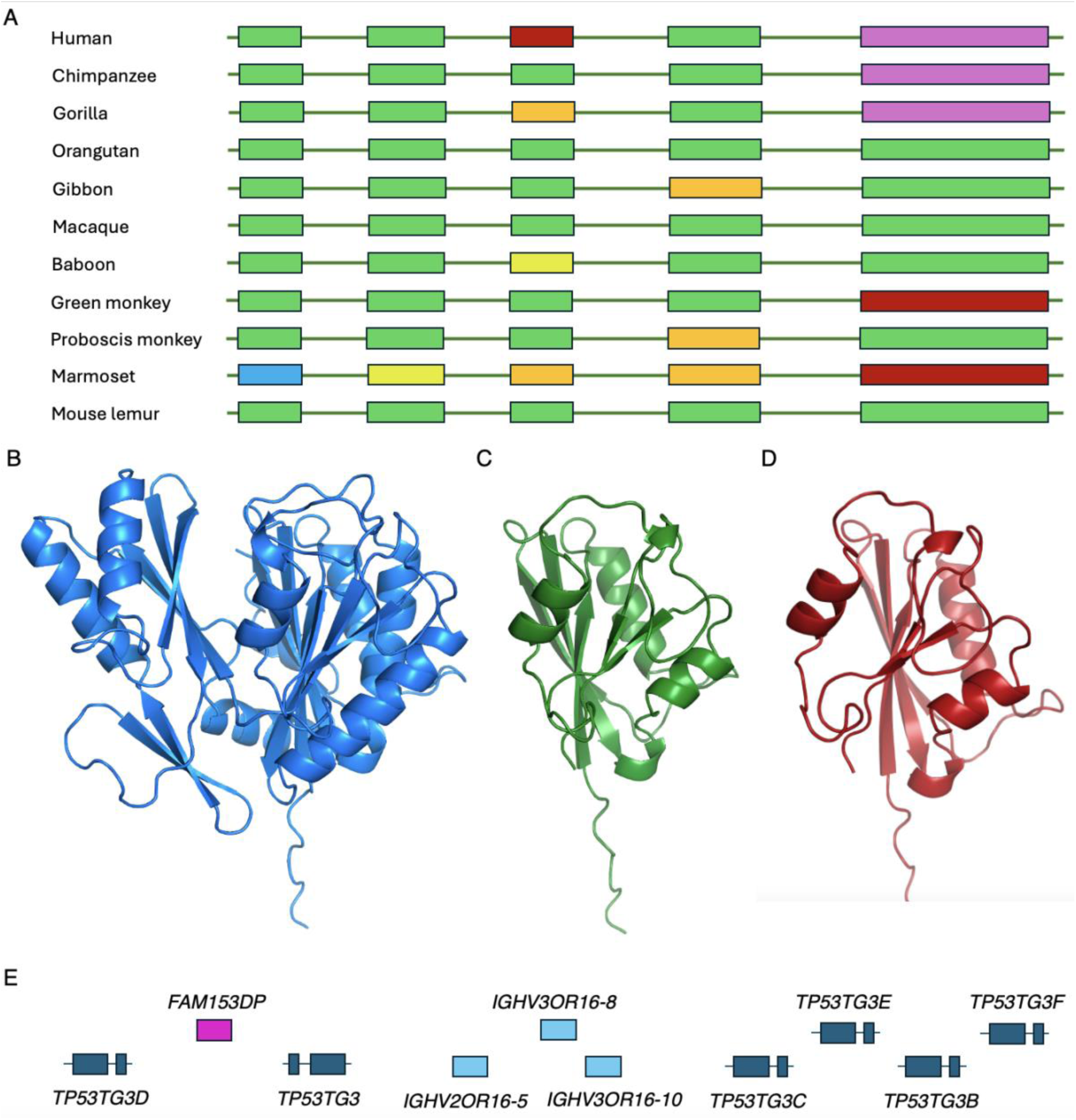
Potential non-coding genes *FTCDNL1* and the *TP53TG3* family. Panel A shows the exon structure of primate *FTCDNL1* genes and pseudogenes. Exons and introns not to scale. Exons missing the ATG are in blue, exons missing splice sites in yellow, exons with frameshift mutations in orange, exons with stop codons in red and missing exons in purple. Panel B shows the AlphaFold predicted structure of cow *FTCDNL1* with both lobes of the structure. Panel C shows the AlphaFold predicted structure of the UniProtKB reference sequence for human *FTCDNL1*. It is missing one lobe of the structure, and the right-hand lobe is missing sections that will not allow it to fold properly. Panel D shows the AlphaFold predicted structure of the MANE Select reference sequence [15] for human *FTCDNL1*. As with the UniProtKB reference sequence, it is also missing a lobe of the structure, and the remaining lobe will fold properly. The APPRIS principal sequence [20] is different again from the two reference sequences shown here, but not any better. Panel E shows the section of chromosome 16 that houses most of the *TP53TG3* orthologues. Exons are not to scale.

Mammalian *FTCDNL1* proteins fold into a single globular domain (figure 6B), but human *FTCDNL1* isoforms are either shortened by the human-specific stop codon in exon 3, or are produced from transcripts that skip exon 3 and terminate in more or less random non-conserved regions downstream of exon 4. The loss of exons 3 and 5 would lead to a radically changed protein structure (figure 6C and 6D). Although *FTCDNL1* seems to be expressed in almost all tissues at the RNA level, there is no evidence that it is translated into protein in either Human Protein Atlas or PeptideAtlas.

Gene *TP53TG3* was discovered in an analysis of a cancer colon cell line a quarter of a century ago [52]. However, no further work has ever been carried out on this gene or any of its paralogues. There are five sequence-identical coding paralogues of *TP53TG3* and two pseudogenes found on the short arm of chromosome 16 (figure 6E). The TP53TG3 genes were all tagged as potential non-coding in the previous analysis [13], but not in this investigation because UniProtKB annotates them as having homology support and this feature is not considered to be potential non-coding in this study. It is not clear which homologous known proteins UniProtKB used as evidence, since there are none.

The TP53TG3 genes are a very recent innovation with conserved ORFs in only chimp and gorilla. Gorillas have one copy of the gene and a pseudogene, and chimp and bonobo have a pseudogene and several TP53TG3 genes. Beyond these species, *TP53TG3* cannot be coding because of the multiple frame shifts and stop codons that break the ORF, and because the splice sites in the various transcripts are not conserved either. Given that, it seems highly unlikely that any of the *TP53TG3* paralogues play any role at all in the cell cycle checkpoint pathway as was originally predicted [52].

There is substantial RNA evidence for *TP53TG3* and its paralogues in testis and epididymis in Human Protein Atlas, and there is also some in cancer cell lines. Despite the substantial RNA evidence in these tissues, there is no peptide evidence in PeptideAtlas. Given the transcript expression, there is no obvious explanation for not detecting peptide evidence in PeptideAtlas. Constructs of *TP53TG3* have a long half-life [52], the protein has multiple detectable tryptic peptides and there are many large-scale testis tissue experiments in PeptideAtlas. Given the lack of conservation and lack of evidence for any protein, it would be remarkable if any of these six genes were protein coding.

## Conclusions

We have carried out a follow up of our in-depth analysis of the coding genes annotated in three main human reference databases [13]. In it we show that there are almost 350 fewer human coding genes annotated across the three sets than in our previous analysis, and that the coding genes agreed upon by all three reference sets has decreased by 179 genes. However, there are still 2,606 genes that the three reference sets cannot agree on.

If read-through genes and immunoglobulin fragments are omitted from the sets, the number of coding genes that the reference databases disagree on falls to slightly over 1,500, and the number of genes agreed upon by all three reference annotators has increased by 248 since we last compared the three annotations in 2018 [13]. Omitting read-through genes from the sets is not just a theoretical exercise, since Ensembl/GENCODE have plans to reclassify all read-through genes that are currently classified as coding. Currently, Ensembl/GENCODE annotates 655 read-through genes as protein coding.

The Ensembl/GENCODE reference set stands out because it only has 114 true coding genes that are not yet coding in RefSeq and UniProtKB. This demonstrates the efficiency of the joint MANE and GIFTS projects that Ensembl/GENCODE shared with RefSeq and UniProtKB and the work that Ensembl/GENCODE have done to help reach a consensus gene set is to be commended. Yet while read-through genes are annotated as coding, the three gene sets are still some way short of this consensus. Read-through genes, along with the unfinished transcript fragments that Ensembl/GENCODE alone annotate, unnecessarily complicate large-scale analyses and many users are unaware of these flaws in the Ensembl/GENCODE annotation.

We also revisited the potential non-coding features that we introduced in our previous analysis. For this analysis we determined potential non-coding features from the non-synonymous to synonymous ratios of genes with these features. With fewer potential non-coding features in this analysis, we predicted fewer potential non-coding genes for the Ensembl/GENCODE reference set. Of the 1,118 Ensembl/GENCODE genes predicted as potential non-coding in this analysis, 843 (75.4%) were outside of the intersection between the three sets.

In this analysis, we also predicted potential non-coding for the other two reference sets. There were 941 UniProtKB genes that were tagged as potential non-coding, more than two thirds (666) were not annotated as coding by either RefSeq or Ensembl/GENCODE or both. For RefSeq we found just 706 potential non-coding genes, but only three of the potential non-coding features could be applied to the RefSeq-unique genes. However, we were able to show that the 683 RefSeq coding genes outside of the intersection were not under purifying selection, so most predicted RefSeq genes that do not agree with the other two annotation sets are unlikely to produce functionally important proteins.

Although we flagged fewer potential non-coding genes than in previous analyses, 1,118 Ensembl/GENCODE genes is still a large number. As is clear from the number of potential non-coding genes that have peptide evidence, not all of these potential non-coding genes will turn out to be non-coding, but 379 were reclassified after the previous analysis [13]. However, as we have shown here, a not insubstantial number of those potential non-coding genes inside the intersection of the three reference sets are not coding genes.

Outside of the intersection between the three reference sets we believe that the vast majority are likely to be non-coding genes or pseudogenes, although there will be exceptions, such as the UniProtKB genes outside the intersection that we recently showed were coding [53, 54] and the recently annotated Ensembl/GENCODE short ORFs with conservation support that seem likely to be coding [4].

Determining whether a gene flagged as potential non-coding is coding or not is not always a simple process. Some genes may have published functional evidence which may complicate what would otherwise be a simple decision [55, 56], while other genes might be supported by evidence from large-scale databases such as Human Peptide Atlas antibodies or PeptideAtlas peptides that need to be contrasted to see if they are reliable. Pseudogenes and coding genes are particularly difficult to disentangle if the ORF is maintained but has no supporting evidence. However, as we have shown here and in an earlier analysis [7], it is not always impossible. Curiously, we have also shown that genes previously thought to be pseudogenes precisely because they had broken ORFs, are in fact coding [53, 54].

One particularly problematic set of genes are those large recently duplicated gene families like the PRAMEF family [57] and the USP17L family [58]. These families have multiple recently duplicated genes and little to no supporting evidence, and it is not clear whether one, none or all of these genes are coding. The SINE-Alu derived NPIPB family [59], for instance, does have some supporting evidence [60], but here it is not obvious whether this evidence supports one or two of the members of the family or all of them. If just one or two of these genes are coding, they are so similar that it is not clear which genes are coding. Currently, all genes in large recently duplicated families are annotated as coding as long as the ORFs are conserved.

Conversely, cancer antigens, a set of genes that previously generated doubts about their coding status are much more clear cut. These ORFs may produce proteins [54] under certain circumstances, but there is increasing evidence that cancers generate much aberrant translation [40]. Cancer antigens should not be annotated as protein coding genes and instead annotators should annotate them as what they are: cancer antigens.

The human reference gene count currently stands at somewhere between 19,950 and 20,485 coding genes depending on which of the three main references is used. We believe that this number is still highly inflated, but the number of coding genes is currently growing rather than decreasing because annotators are adding short ORFs [14]. Often these small ORFs overlap UTR or coding exons and most of them have clear protein coding conservation. The annotation of short ORFs as coding genes has been much heralded [12] and seems to be in its early stages.

This work is an analysis of the reference coding gene set from the GRCh38 assembly of the human genome. The annotation of both the new CHM13-T2T assembly [61,62] and the human pangenome [63] have the potential to add to the gene numbers here. The CHM13 assembly completed the remaining 8% of the human genome and unearthed 140 novel genes with protein coding gene pedigree. We found supporting evidence for two novel protein coding genes in the CHM13-T2T assembly [7], but most of these genes were from large, recently duplicated families. For example, 31 of the new genes are duplications of *TAF11L5* and 34 are most similar to the USP17L genes [61]. The new CHM13-T2T assembly is a hugely important achievement for many reasons, but it seems unlikely that it will add more than a handful of new coding genes to those already known.

We believe that the three main reference databases still annotate as protein coding close to 2,000 genes that never produce proteins or that only produce proteins in abnormal conditions such as cancers. These genes should be reclassified because their presence in the reference set of coding genes can only complicate large-scale biomedical experiments. Determining a final agreed upon set of reference coding genes is of fundamental importance to researchers and we hope that this work will inspire the three main reference databases and perhaps others [64] to continue the work towards a final agreed set of *bona fide* coding genes.

## Acknowledgements

This work was funded by the National Human Genome Research Institute of the National Institutes of Health (grant number U41 HG007234).

## Notes

### Competing Interest Statement

The authors are part of the GENCODE consortium

