## Supplementary for "More than 2,500 coding genes in the human reference gene set still have unsettled status"

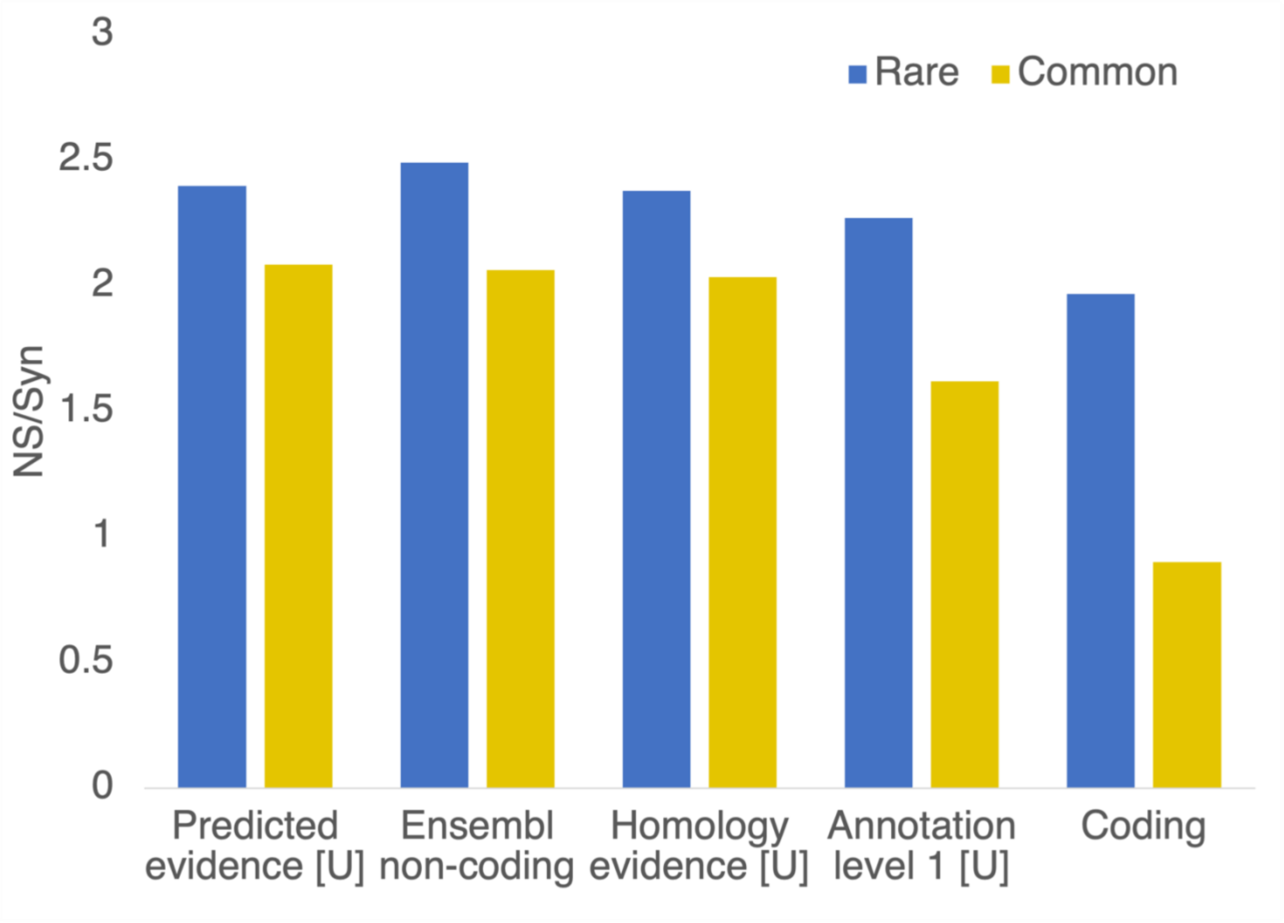


**Supplementary Figure 1**
The NS/Syn ratios calculated for potential non-coding features excluded from the analysis. Features marked with a “U” are annotator evidence codes that come directly from the UniProtKB database. Ensembl non-coding genes are those marked as non-coding, non-functiona or pseudogene. Coding gene NS/Syn ratios (from genes under purifying selection) are included as a control.

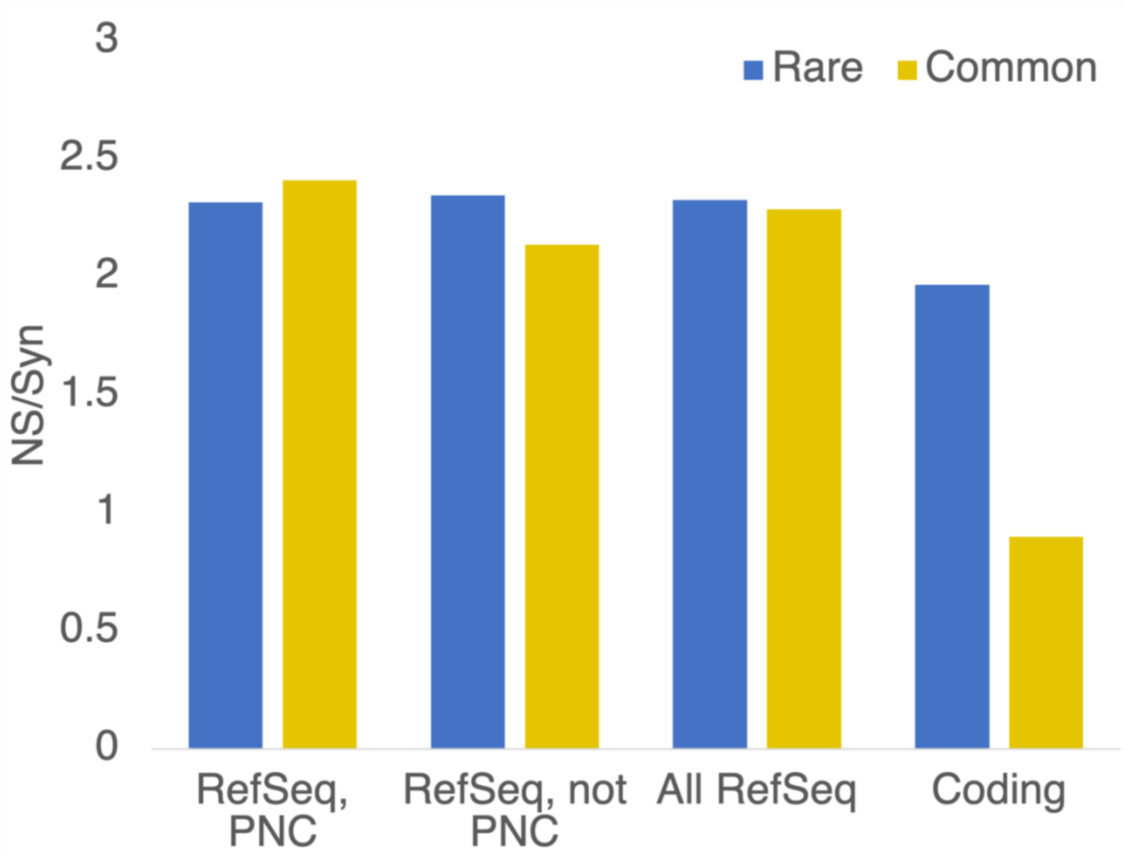


**Supplementary Figure 2**
The NS/Syn ratios calculated for predicted RefSeq genes not in the intersection. “RefSeq, PNC” are those RefSeq coding genes not in the intersection of the three sets that are tagged with one of the three applicable potential non-coding features. “RefSeq, not PNC” are RefSeq coding genes not in the intersection that are not tagged with potential non-coding features. “All RefSeq” are RefSeq coding genes not in the intersection. Coding gene NS/Syn ratios (from genes under purifying selection) are included as a control.

```
HIG2DB HUMAN       MA-----------TLGFVTPEAPFESSKPPIFEGLSPTVY-S-NPEGFKEKFLRKTRENP
MARMOSET           MA-----------TPGPVTPEASLEPSKPPVIEGFSPTLY-S-KPESFKEKFIRKVRENP
MACAQUE            MA-----------TPGPVIPEVPFEPSKPPVIEGLSPTVY-R-DPETFKEKFLRKTRENP
HIG2DA HUMAN       MA-----------TPGPVIPEVPFEPSKPPVIEGLSPTVY-R-NPESFKEKFVRKTRENP
RAT                MA-----------APGPVSPEAPFDPSKAPVIEGFSPTVY-T-NPEGFKEKFIRKTRENP
MOUSE              MA-----------APGPVSPEAPFDPSKPPVIEGFSPTVY-S-NPEGFKEKFIRKTRENP
HAMSTER            MA-----------APRPVSSEAPFDPSQTPVIEGFTPTVY-S-NPESFKDKFIRKTRENP
BAT                MT-----------TPGPVTPGTPFEPSQPPVVEGISPSVY-S-TTESFKEKFLRKTRENP
GUINEA PIG         MA-----------TPGPVTPEAPFKPSQPPVIEGFNPSVH-I-HQEGFKEKFLRKTRENP
PIG                MA-----------TPGPATPEAPFEPSHPPVIEGFSPTVY-S-TSESFKKKFIRKTRENP
TARSIER            MA-----------TPGPVTPEVPFEPSQPPVIEGFSPSVY-G-NPESFKEKFLRKTRENP
CAT                MA-----------VPGPVTPEAPFEPSQPPVIEGFSPSVY-S-TPESFKEKFLRKTRENP
DOG                MA-----------APGPVTPGAPFEPSQPPVIEGFSPSVY-S-PPESFKEKFLRKTRENP
FOX                MA-----------APGPVTPGAPFEPSQPPVIEGFSPSIY-S-TPESFKEKFLRKTRENP
POLAR BEAR         MA-----------APGPVTPGAPFEPSQPPVIEGFSPSIY-N-TQESFKEKFLRKTRENP
SEAL               MA-----------APGPVTPGAPFEPSQPPVIEGFSPSIY-S-TQESFKEKFLRKTRENP
DOLPHIN            MA-----------TPGPVTPEAPFEPSQPPVIEGFSPSVY-S-TSESFKEKFLRKTRENP
SHEEP              ME-----------TPGRVTPEAPFEPSQPPVIEGFSPSVY-S-TSEGFKEKFIRKTRENP
COW                ME-----------TPGRVTPEAPFEPSQPPVIEGFSPSVY-S-TSESFKEKFIRKTRENP
KILLFISH           MAATTRPSEPESHVKEQPPGALPFVLSQPPVIEGFRQSPK-V-KDETFKEKFIRKTKENP
SNAILFISH          MA-GATAVSELTPAAKSPAAFAPFDFSQPPDIEGFSRLPA-A-RDETFKEKFLRKTKENP
TROPICAL FROG      --------------------------MAHPEIEGFTPSST-Y-GDEGFKSKFIRKVKENP
```

```
AFRICAN FROG      --------------------------MAQPEIEGFTPS-T-Y-TSEGFKGKFIRKVKENP
SHARK             ---------------MAAPVREEVSLSKPPVIEGFHPAPR-P-REESFRDKFIRKTSENP
COD               MAAASTPAVVNSPI-TQQSGGMPFDFSKPPVIEGFTPVSR-R-KDETFKEKLLRKTKENP
TROUT             MAAATTPVVPEQSASTT--HLPMLDISKPPVIDGFTPLSR-P-REETFQEKFMRKSKENP
CAVE FISH         MAVASSAVQEERAGRTALPGAAAFNPGSPPVIEGFTPLPR-Q-KEESFKEKFLRKTKENP
ZEBRAFISH         MATAAAPVSPDQPGKSA-SPPVLLDLSQPPVIEGFSPTSR-T-REEGFKDKFIRKTKENP
CATFISH           MAAASSRVEQASPVKTAPPIAAGFDLSDPPVIDGFTPLSR-P-REEGFKDKFIRKTKENP
HERRING           MAASPTVVSQDQAANAASQGPIPFDISKPPIIEGFTPLPR-H-REEGFKDKFIRKTKENP
MILKFISH          MAAASTPVERDQVVKTPQSGPVPFDISKPPVIEGFTPLPR-A-KEEGFKDKFIRKTKENP
CHICKEN           ---------------MAAGPPPPLEPIPLPVY-----------RDEGFADKFRRKTRENP
ZEBRA FINCH       ---------------MAAGPPPPLEPSPLPTF-----------PEEGFTEKFVRKTRENP
BARN SWALLOW      ---------------MAAGPPPPLDPIPLPTF-----------TEEGFAEKFLRKTRENP
WALL LIZARD       ---------------MAQSAPPPFDPNNPPLIEGFTPTAY-H-PEEGFGDKFRRKTRENP
ANOLE LIZARD      ---------------MTQAPPPPFDPSRPPLIEGFQPGAF-QRREEGFADKFLRKTRENP
                                               *               * *  *: **  ***

HIG2DB HUMAN      VVPIGFLCTAAVLTNGLYCFHQGNSQCSRLMMHTQIAAQGFTIAAILLGLAATAMKSPP-
MARMOSET          MVPIGCLATATALGYGLYCFHKGHSRRSQLMMRTRIAAQGFTIAAILVGLGVTSMKSRP-
MACAQUE           VVPIGCLATVAALTYGVYSFYRGDSRRSQLMMRTRIAAQGFTVTALLLGLAVTAMKSRP-
HIG2DA HUMAN      VVPIGCLATAAALTYGLYSFHRGNSQRSQLMMRTRIAAQGFTVAAILLGLAVTAMKSRP-
RAT               MVPIGCLGTAAALTYGLYCFHRGQSHRSQLMMRTRIAAQGFTVVAILLGLAASTMKSRS-
MOUSE             MVPIGCLGTAAALTYGLYCFHRGQSHRSQLMMRTRIAAQGFTVVAILLGLAASAMKSQA-
HAMSTER           MVPIGCLGTAAALSYGLYCFHRGQSHRSQIMMRTRIAAQGFTVVAILLGLAASAMKSRS-
BAT               MVPLGCLSTAAALTYGLYCFHRGQSQRSQLMMRTRIAAQGFTIVAILVGLAASALKSRP-
GUINEA PIG        MVPIGCLGTAAALTYGLYCFHQGHSQRSQFMMRTRIAAQGFTVAAILLGLAASAMKSRS-
PIG               MVPIGCLGTASALTYGLYCFHRGQSQRSQLMMRTRIAAQGFTIVVILVGLAASTMRSRP-
TARSIER           MVPIGCLGTAAALTYGLYCFHRGHSQRSQLMMRTRIAAQGFTVAAILLGLAASALKSRP-
CAT               MVPIGCLGTAAALTYGLYCFHRGQSHRSQLMMRTRIAAQGFTVAAILLGLAASAMRSRS-
DOG               MVPVGCLGTAAALTYGLYCFHRGQSHRSQLMMRTRIAAQGFTVAAILLGLAASAMKSRS-
FOX               MVPIGCLGTAAALTYGLYCFHRGQSHRSQLMMRTRIAAQGFTVAAILLGLAASAMKSRS-
POLAR BEAR        MVPIGCLGTAAALTYGLYCFHRGQSHRSQLMMRTRIAAQGFTVAAILLGLAASAMKSRS-
SEAL              MVPIGCLGTAAALTYGLYCFHRGQSHRSQLMMRTRIAAQGFTVAAVLLGLAASAMKSRS-
DOLPHIN           MVPIGCLGTAAALTYGLYCFHRGQSQRSQLMMRTRIAAQGFTVVAILMGLAASTMKSRP-
SHEEP             LVPIGCLGTAAALTYGLYCFHRGQSQRSQLMMRTRIAAQGFTIVAILVGLAASTLKSRP-
COW               LVPIGCLGTAAALTYGLYCFHRGQSQRSQLMMRTRIAAQGFTIVAILVGLAASTLKSRP-
KILLFISH          FVPIGCLGTTGMLMYGLRSFHQGKTKQSQMFMRGRIFAQGFTVVAIIVGIFATALKPKQ-
SNAILFISH         FVPIGCLGTAGALVYGLRAFNQGKTRQSQLMMRGRIFAQGFTVVAIIAGVFITAMKPKQ-
TROPICAL FROG     FVPIGCLATAGALTYGLISFKQGKTQQSQLLMRTRILAQGFTVAAIMFGVVMTAMKPRIT
AFRICAN FROG      FVPIGCLATAGALTYGLISFKQGKTRQSQLLMRTRILAQGFTVAAIMVGVVMTALKPSET
SHARK             FVPLGMLGTAGALTYGLIAFNHGKTRHSQLSMRARIFAQGFTIVAIVVGVVATTLKPK-
COD               FVPIGCLGTAGALAYGLRAFHQGKTRQSQMLMRGRIFAQGFTVFAIIFGVVATQLKPKQ-
TROUT             FVPIGCLGTAGALMYGLRAFKQGKTRQSQLLMRGRIFAQGFTVVAIIFGVFTTALKKD--
CAVE FISH         FVPIGCLGTAGALTYGLRAFKHGKTHQSQMLMRTRIFAQGFTVIAIIVGVAATALKSKQ-
ZEBRAFISH         FVPIGCLGTAGALIYGLGAFKQGKTRQSQLLMRTRIFAQGFTVVAIIVGVAATALKAKP-
CATFISH           FVPIGCLGTAGALIYGLRAFKQGKTRQSQLLMRTRIFAQGFTVVAIIVGVAAAALKPRQ-
HERRING           FVPIGCLGTAGALIYGLRAFKMGKTRQSQLSMRMRIFAQGFTVVAIIVGVASTALKPKQ-
MILKFISH          FVPIGCLGTAGALIYGLSAFRQGKTRQSQLLMRARIFAQGFTVVAIIVGVATTALKSK-
CHICKEN           LVPLGCLCTLGVLTYGLISFKRGNTRHSQLMMRARVVAQGFTVAALLGGMVATALRARS-
ZEBRA FINCH       LVPLGCLCTVSVLVYGIICFKRGQTRRSQLMMRARVIAQGCTFAALLGGMVATALKSRQ-
BARN SWALLOW      MVPLGCLCTVGVLAYGVICFKKGNTRRSQLMMRARVVAQGFTIASVVGGMMATAIRSRQ-
WALL LIZARD       LVPIGCLGTAGVLAYGLICFKKGNTLQSQRMMRARVLAQGFTVAAILVGVVVASMKPKK-
ANOLE LIZARD      LVPVGCLGTAGVLTYGLICFKRGNTHQSQIMMRARILAQGFTVAALVVGVVVTALKPKK-
                  .**:* * *   * **: .*  *.   **  ** *: *** *.  :: *:     :
```

**Supplementary Figure 3. Alignment of human *HIGD2B* with vertebrate *HIGD2A* homologues**
Alignment of human *HIG2DB* with 35 vertebrate *HIGD2A* paralogues. *HIGD2A* sequence is shown in bold with the 29 residues that are completely conserved across the 35 *HIGD2A* in blue. *HIGD2B* sequence is shown in bold with residues that are different from completely conserved *HIGD2A* residues in red, residues that are radically different from highly conserved *HIGD2A* residues in orange and other residues that are radically different in green.
